# Risk of complications among diabetics self-reporting oral health status in Canada: A population-based cohort study

**DOI:** 10.1101/652529

**Authors:** Kamini Kaura Parbhakar, Laura C. Rosella, Sonica Singhal, Carlos R. Quiñonez

**Affiliations:** Department of Dental Public Health, Faculty of Dentistry, University of Toronto, Toronto, Ontario, Canada; Division of Epidemiology, Dalla Lana School of Public Health, University of Toronto, Toronto, Ontario, Canada; Institute for Clinical Evaluative Sciences, Toronto, Ontario, Canada; Public Health Ontario Toronto, Ontario, Canada

## Abstract

**Background:** Periodontitis has persistently been associated with diabetes and poor health outcomes. While clear associations have been identified for the diabetes–oral health link, less is known about the implications of poor oral health on incident complications of diabetes. This study sought to investigate the risk of diabetes complications associated with self-reported “poor to fair” and “good to excellent” oral health status among diabetics living in Ontario, Canada.

**Methods:** This cohort study was undertaken of diabetics from the Canadian Community Health Survey (2003 and 2007-8). Self-reported oral health was linked to electronic health records at the Institute for Clinical Evaluative Sciences. Participants under the age of 40, missing self-reported oral health and those who could not be identified in linked databases were excluded (N=5,183). A series of Cox Proportional hazard models were constructed to determine the risk of diabetes complications. Participants who did not experience any diabetes complication were censored at time of death or at the study termination date (March 31, 2016). Models were adjusted for age and sex, followed by social characteristics and behavioural factors.

**Results:** Diabetes complications differed by self-reported oral health. For those reporting “poor to fair” oral health, the hazard of a diabetes complication was 30% greater (HR 1.29 95%CI 1.03, 1.61) than those reporting “good to excellent” oral health.

**Conclusions:** Our findings indicate that oral health status is associated with increased risk for complications among diabetics, after adjusting for a wide range of confounders. Examining oral health and the risk for diabetes complications from a broader perspective including socio-behavioural and biological pathways is principal for informing policies and interventions that aim to mitigate the burdens of poor systemic health.

## Introduction

Diabetes is one of the most prevalent chronic conditions and the 6^th^ leading cause of mortality in Canada (1,2). The epidemic continues to grow as an increasing number of Canadians are living with diabetes and prediabetes, and is expected to impact 13.3 million individuals by 2029 (3). And although an abundance of research and literature has focused on diabetes control and management of complications, including cardiovascular diseases, renal disease, retinopathy and amputations; less attention has been given to periodontal disease, the 6^th^ most common complication of diabetes (4).

Periodontal disease is a dysbiotic inflammatory condition that destroys the bone and connective tissue supporting teeth; diabetes is a condition affecting glucose homeostasis. Recent meta-analyses have reported that mild to moderate periodontal disease affects a majority of adults, and approximately 5%-20% of any population suffers from the severest form (5–7). Mild periodontal disease is characterized as an acute form of the condition and severe periodontal disease occurs as the result of a chronic state of disease. 6% of Canadian adults have severe periodontal disease (8). However, 3 times the number of diabetics have a combined prevalence of moderate to severe periodontal disease, making this 6^th^ complication an important concern (9)

Both diabetes and periodontal disease facilitate inflammatory pathology at local and distant sites and many studies have presented evidence regarding the complex relationship between these two chronic conditions (7,10). The low grade immune response that develops from periodontal infection can lead to systemic levels of inflammation, exaggerating the immune-inflammatory response that is instrumental to insulin resistance, thereby increasing the severity of diabetes (11–15). It is this host-mediated response which drives the bidirectional relationship between these chronic diseases.

The bidirectional link between periodontal disease and diabetes has received considerable global attention. Yet, the implications of periodontal disease and oral health on diabetics in Canada is unknown. A 2015 systematic review concluded that periodontal therapy reduced blood sugar levels following care, thereby improving metabolic control among diabetics (4). An earlier cohort study reported that study participants who received periodontal therapy experienced a reduction in the risk of diabetes complications and medical costs (16). Similar studies conducted from 2008-2016 reported reduced pharmaceutical costs, increased immediate and long-term medical cost savings, as well as reduced hospital admissions and physician visits among individuals receiving periodontal treatment (17–19). As both the prevalence of diabetes and its associated indirect and direct costs continue to grow worldwide (20,21), periodontal therapy may be a solution to a lack of metabolic control among diabetics with the severest of conditions; population level interventions addressing oral health and diabetes could also improve health outcomes for Canadians.

However, there is a paucity of population level evidence on the epidemiological association between oral health and diabetes complications in Canada. Much of the recent literature has been conducted in clinical settings and in cohorts that were followed for varying timeframes rendering results of low to medium quality (4). Moreover, systematic reviews have reported mixed conclusions, warranting the need for further research. This study thus aims to identify the risk of diabetes complications among a cohort of diabetics self-reporting oral health status, in Ontario, Canada’s most populated province.

## Research Design and Methods

### Study Population

This cohort study was designed with the objective of assessing the risk of diabetes complications among diabetics self-reporting oral health status. The base cohort comprised of Ontario residents who participated in the 2003 and 2007-08 Canadian Community Health Survey (CCHS). The CCHS is a cross-sectional survey annually administered by Statistics Canada and collects self-reported health data. Briefly, the CCHS utilizes a multi-stage, stratified, clustered-probability survey sampling design that is administered to 98% of the Canadian population that is 12 years of age or older, excluding those living on reserves, individuals residing within institutions, and full-time members of the Canadian Armed Forces (22). Details regarding the CCHS survey methodology are documented elsewhere (22).

As all residents of the province of Ontario are covered by a single payer insurance system known as the Ontario Health Insurance Plan (OHIP), health system encounters can be tracked. With de-identified OHIP health card numbers, health encounters are recorded in administrative health data and held at the Institute for Clinical Evaluative Sciences (ICES). The Ontario Diabetes Database (ODD) is an ICES-derived disease registry used to identify all physician-diagnosed cases of diabetes in Ontario (23). The ODD uses the diagnostic criteria of two physician service claims recorded in OHIP or one hospital discharge related to diabetes within a two-year period to identify incident diabetes cases. The ODD has been validated with a sensitivity of 86% and a specificity of 97% for classifying individuals with and without type 2 diabetes (23).

The final study sample was restricted to CCHS participants (2003 or 2007-8), over the age of 40 years at the interview date, with a diabetes diagnosis from the ODD. Individuals who did not participate in the oral health component of the CCHS or were OHIP-ineligible for the entire observation window were excluded. The final cohort consisted of 5,183 individuals, representing a weighted sample of 1.31 million Ontario residents.

### Oral Health

The oral health content of CCHS cycles 2003 and 2007-08 were selected to ensure the availability of oral health linked electronic medical records. Cycles were combined using a pooled approach to increase the sample size and provide greater statistical power (24). Self-reported oral health status is the exposure variable and was used as a proxy for periodontal condition. It was assessed through the question “would you say the health of your teeth and mouth is: excellent, very good, good, fair, and poor?” Based on the distribution of respondents in these categories, the variable was dichotomized into “good to excellent” versus “poor to fair” oral health groups. Further description of oral health content of the CCHS is described elsewhere (22).

### Diabetes complications

CCHS respondents, with a diabetes diagnosis, were followed prospectively from the survey interview date until March 31, 2016. Only the first diabetes complication following the interview date was captured from electronic medical records and those who did not experience a diabetes complication or died before the end of the follow-up period were censored. Participants were dichotomized according to those who experienced a complication and those who did not. Complications were defined by select International Classification of Disease (ICD) codes previously used in the literature (25). Complications included the following: hyper- and hypoglycemia, myocardial infarction, stroke, skin infection, amputation, kidney failure, dialysis and retinopathy.

### Baseline Covariates

At the CCHS interview date, survey participants also reported demographic characteristics, health behaviours as well as medical histories. The following covariates from the CCHS were included in our analysis: age, gender, income, education, immigrant status, race, physical activity, smoking status, alcohol consumption, dental visits, BMI, comorbidity, stress, and self-reported overall health.

Physical activity was measured using an index derived from self-reported physical activities in the past 3 months prior to interview date (26). Type of smoker was derived using lifetime cigarette consumption and was categorized as current smoker, former smoker and never smoked (26). On the other hand, alcohol consumption was characterized as regular, occasional, former or never had a drink in the past 12 months (26). Stress was defined by the amount of stress perceived by study participants on most days and was dichotomized into “perceiving stress” and not “perceiving stress”. Comorbidity was assessed by any self-reported chronic disease diagnosed by a physician prior to interview date, other than diabetes. Participants were dichotomized into groups without any comorbidity or those with at least one of the following at interview date: arthritis, COPD, heart disease or stroke. Due to the distribution of individuals’ ethnicity, this variable was dichotomized into “white Caucasian” and “ethnic minority” categories, which was comprised of Black, South Asian, Latin American, Indigenous and other ethnicities.

Other covariates extracted from electronic medical records include rural-urban dwelling, the duration of diabetes prior to the interview date as well as medical care received prior to the first complication experienced. The Rurality Index of Ontario (RIO) was used to distinguish participants residing in rural areas versus urban areas. RIO scores of 0-39 were considered urban dwelling and scores greater than 40 were considered rural dwelling (27). The duration of diabetes prior to interview date was captured from the ODD. And the type of medical care was assessed with OHIP codes for general practitioner (GP) and/or specialist visits. Medical care could be provided by a GP, specialist, or by both a GP and specialist prior to the diabetes complication.

### Statistical analysis

Descriptive statistics were calculated by oral health status and included means, standard deviations and distributions of demographic and health behaviour variables. Baseline characteristics were compared among participants self-reporting “good to excellent” and “poor to fair” oral health status using t-tests for continuous variables and chi-squared tests for categorical variables. Bivariate analysis was then conducted for all variables by the dichotomous outcome. A p-value of 0.25 was used as a cut-off point in the bivariate analysis in order to qualify variables for a forward building model (28).

Cox proportional hazards models were built using person-days as the time-to-event for the first diabetes event. Parsimonious models were built and multivariable hazard ratios (HRs) with confidence intervals (CIs) were estimated for diabetes events associated with self-reported oral health. The proportional hazard assumption was met by using the log rank test. The survival curves from the log rank test of diabetes events by self-reported oral health status are shown in figure 1. Individuals self-reporting “good to excellent” oral health represented the reference category.

**Figure 1:**
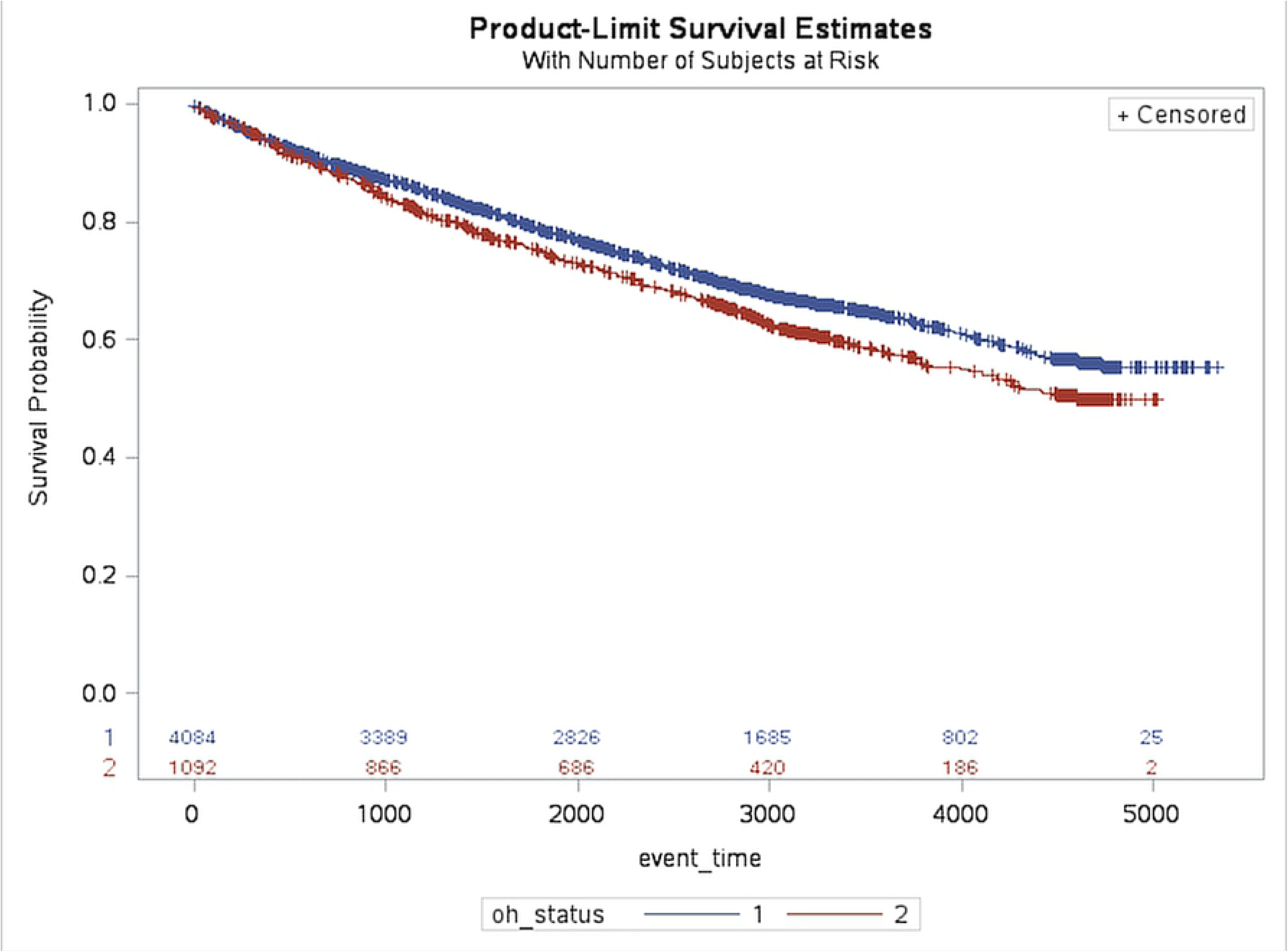
Log-rank test for probabilities of diabetes complications among study participants self-reporting “good to excellent” (oh_status=1) and “poor to fair” (oh_status =2) oral health status over event time in days. (n= 5,183; N= 1,308,911)

Four primary models were constructed to test the association between self-reported oral health and diabetes complications. Following a base crude model (model 1), model 2 adjusted for age and sex. This was followed by additional adjustments for social demographic factors including income, education, ethnicity, marital status, immigrant status and RIO scores (model 3). A final adjustment in model 4 was conducted for behavioural and other influencing factors including BMI, activity index, and alcohol consumption, smoking behaviour, comorbidity, sense of community belonging and dental visits.

Bootstrapping sample weights provided by Statistics Canada were applied to the analyses to adjust for the complex survey design of the CCHS and to generate inferable estimates for the Ontario population. All statistical analyses were performed using SAS version 9.4 (29). PROC SURVEY FREQ, PROC SURVEY MEANS AND PROC SURVEY PHREG were used in SAS. All CCHS respondents provided consent for Registered Persons Database (RPDB) linkage with electronic medical records during CCHS administration. Ethical approval for this study was obtained from the University of Toronto Health Sciences Research Ethics Board followed by subsequent ICES approval for data creation and access (protocol reference #34553).

## Results

The baseline characteristics of study participants according to self-reported oral health status have been shown in table 1. Participants reporting “poor to fair” oral health status were found to be a mean age of 62 years, male and of white ethnicity. In comparison to participants self-reporting “good to excellent” oral health status, those with “poor to fair” oral health were predominantly from the lowest income quintile, were less educated, reported comorbidity prior to interview date and had a higher mean BMI. They also reported “poor to fair” overall health and mental health status, as well as a lower sense of community belonging. They were current smokers, living inactive lifestyles, and making fewer dental visits than those reporting “good to excellent” oral health. During the observation window, 35% of all participants had a diabetes complication following the interview date (table 2). 38% of those reporting “poor to fair” oral health status had a diabetes complication in comparison to 34% of those reporting “good to excellent” oral health (table 2).

**Table 1:**
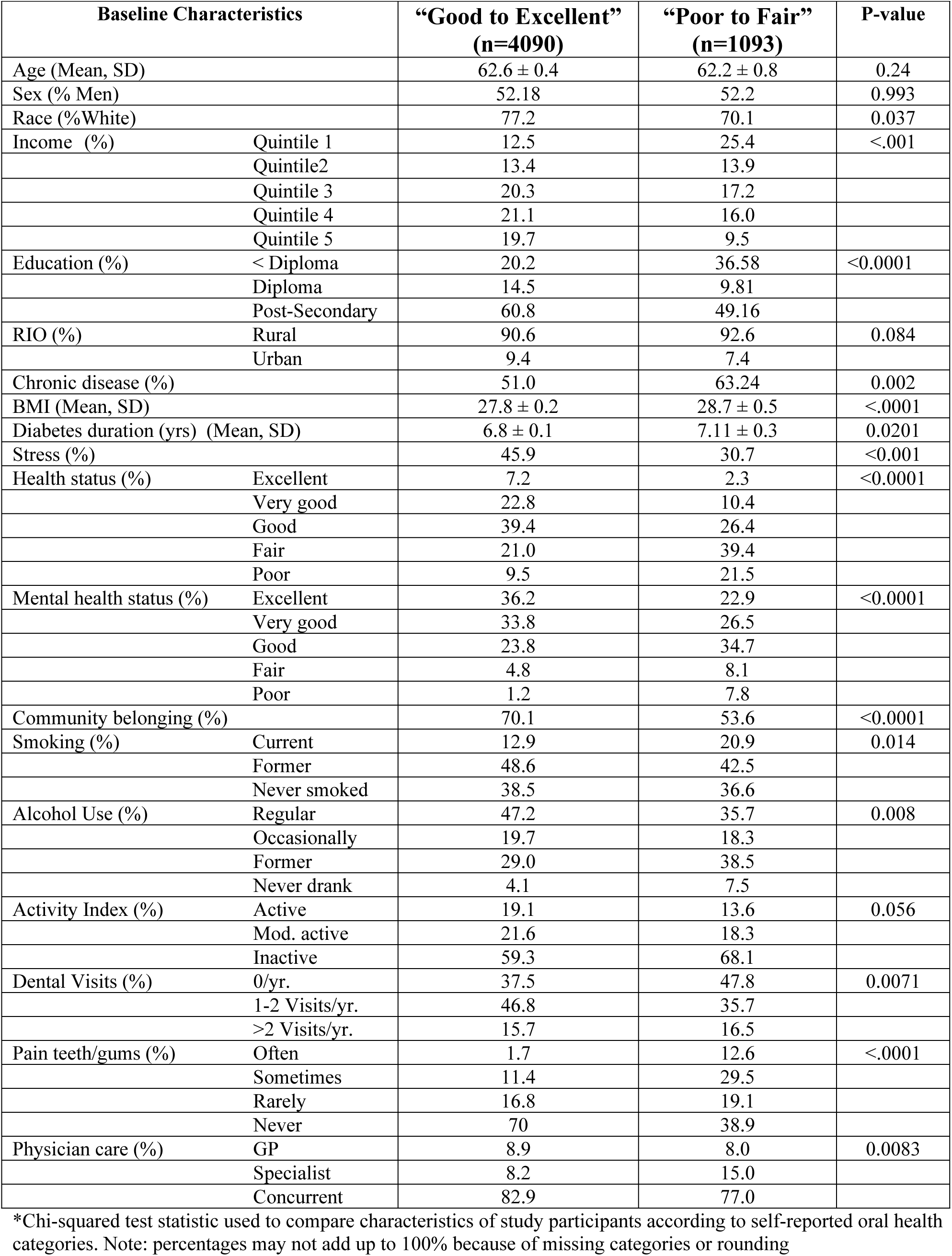
Baseline weighted characteristics of CCHS survey participants according to self-reported oral health status (n= 5,183; N= 1,308,911)

| Baseline Characteristics |  | “Good to Excellent”<br>(n=4090) | “Poor to Fair”<br>(n=1093) | P-value |
| --- | --- | --- | --- | --- |
| Age (Mean, SD) |  | 62.6 ± 0.4 | 62.2 ± 0.8 | 0.24 |
| Sex (% Men) |  | 52.18 | 52.2 | 0.993 |
| Race (%White) |  | 77.2 | 70.1 | 0.037 |
| Income (%) | Quintile 1 | 12.5 | 25.4 | <.001 |
|  | Quintile2 | 13.4 | 13.9 |  |
|  | Quintile 3 | 20.3 | 17.2 |  |
|  | Quintile 4 | 21.1 | 16.0 |  |
|  | Quintile 5 | 19.7 | 9.5 |  |
| Education (%) | < Diploma | 20.2 | 36.58 | <0.0001 |
|  | Diploma | 14.5 | 9.81 |  |
|  | Post-Secondary | 60.8 | 49.16 |  |
| RIO (%) | Rural | 90.6 | 92.6 | 0.084 |
|  | Urban | 9.4 | 7.4 |  |
| Chronic disease (%) |  | 51.0 | 63.24 | 0.002 |
| BMI (Mean, SD) |  | 27.8 ± 0.2 | 28.7 ± 0.5 | <.0001 |
| Diabetes duration (yrs) (Mean, SD) |  | 6.8 ± 0.1 | 7.11 ± 0.3 | 0.0201 |
| Stress (%) |  | 45.9 | 30.7 | <0.001 |
| Health status (%) | Excellent | 7.2 | 2.3 | <0.0001 |
|  | Very good | 22.8 | 10.4 |  |
|  | Good | 39.4 | 26.4 |  |
|  | Fair | 21.0 | 39.4 |  |
|  | Poor | 9.5 | 21.5 |  |
| Mental health status (%) | Excellent | 36.2 | 22.9 | <0.0001 |
|  | Very good | 33.8 | 26.5 |  |
|  | Good | 23.8 | 34.7 |  |
|  | Fair | 4.8 | 8.1 |  |
|  | Poor | 1.2 | 7.8 |  |
| Community belonging (%) |  | 70.1 | 53.6 | <0.0001 |
| Smoking (%) | Current | 12.9 | 20.9 | 0.014 |
|  | Former | 48.6 | 42.5 |  |
|  | Never smoked | 38.5 | 36.6 |  |
| Alcohol Use (%) | Regular | 47.2 | 35.7 | 0.008 |
|  | Occasionally | 19.7 | 18.3 |  |
|  | Former | 29.0 | 38.5 |  |
|  | Never drank | 4.1 | 7.5 |  |
| Activity Index (%) | Active | 19.1 | 13.6 | 0.056 |
|  | Mod. active | 21.6 | 18.3 |  |
|  | Inactive | 59.3 | 68.1 |  |
| Dental Visits (%) | 0/yr. | 37.5 | 47.8 | 0.0071 |
|  | 1-2 Visits/yr. | 46.8 | 35.7 |  |
|  | >2 Visits/yr. | 15.7 | 16.5 |  |
| Pain teeth/gums (%) | Often | 1.7 | 12.6 | <.0001 |
|  | Sometimes | 11.4 | 29.5 |  |
|  | Rarely | 16.8 | 19.1 |  |
|  | Never | 70 | 38.9 |  |
| Physician care (%) | GP | 8.9 | 8.0 | 0.0083 |
|  | Specialist | 8.2 | 15.0 |  |
|  | Concurrent | 82.9 | 77.0 |  |

\*Chi-squared test statistic used to compare characteristics of study participants according to self-reported oral health categories. Note: percentages may not add up to 100% because of missing categories or rounding

**Table 2:** Diabetes complications experienced by participants self-reporting their oral health status (n= 5,183; N= 1,308,911)

| Oral Health Status | Event Type |  | Total |
| --- | --- | --- | --- |
|  | No complication | Complication |  |
| “good to excellent” | 2693 (66%) | 1397 (34%) | 4090 |
| “fair to poor” | 674 (62%) | 419 (38%) | 1093 |
| Total | 3367 | 1816 | 5183 |
| p=0.008 |  |  |  |

\*percentages calculated are row percentages

The probability of a diabetes complication by self-reported oral health status is reported in Figure 1. As shown, individuals reporting “poor to fair” oral health have a slightly lower survival probability over time, in comparison to individuals reporting “good to excellent” oral health. The proportional hazard assumption was satisfied for Cox Proportional Hazard analysis, as the log rank test resulted in a p-value of <0.01, refuting the null hypothesis that there is no difference between study participants self-reporting “good to excellent” and “poor to fair” oral health status.

The hazard ratios, as shown in Table 3, depict the difference in risk of diabetes complications among participants self-reporting oral health status. The age and sex adjusted risk of a diabetes complications among participants reporting “poor to fair” oral health status is 49% greater than those reporting “good to excellent” oral health status [HR = 1.49, 95%CI = 1.16-1.92]. Additional adjustment for social demographic factors including income, education, race, and rural urban dwelling, only slightly attenuated the association between diabetes complication events and oral health status [HR=1.48, 95%CI=1.18-1.85]. Further adjusting for behavioural factors resulted in the largest reduction in risk of diabetes complications. Participants reporting “poor to fair” oral health status had a 29% greater risk of a diabetes complication event, as compared to those reporting “good to excellent” oral health status [HR = 1.29, 95%CI = 1.03-1.61] in the fully adjusted model. All hazard ratios were significant at p<0.05.

**Table 3:** Multivariable hazard ratios for diabetes complication risk by self-reported oral health status (n=5183)

| Oral Health Status | Model 1<sup>a</sup> | Model 2<sup>b</sup> | Model 3<sup>c</sup> | Model 4<sup>d</sup> |
| --- | --- | --- | --- | --- |
| Good – Excellent<br>(Reference Group) | 1.00 | 1.00 | 1.00 | 1.00 |
| Poor - Fair<br>95% CI | 1.47<br>(1.15, 1.87) | 1.49<br>(1.16,1.92) | 1.48<br>(1.19, 1.85) | 1.29<br>(1.03, 1.61) |

a. Model 1 is the crude model
b. Model 2 adjusted for age, sex
c. Model 3 adjusted for age, sex, income, education, ethnicity, marital status, immigrant status, RIO
d. Model 4 adjusted for age, sex, income, education, ethnicity, marital status, immigrant status, RIO, BMI, activity index, alcohol use, smoking, comorbidity, dental visits

## Discussion and Conclusion

To our knowledge, this is the first study to investigate the association between oral health status and diabetes complications in a prospective cohort in Canada. In general, our findings indicate that self-reported oral health status is associated with an increased risk of diabetes complications after adjusting for a wide range of confounders. This aligns with current evidence regarding the association between periodontal disease and the extent of its severity, with diabetes complications including retinopathy, neuropathy, chronic kidney disease and cardiovascular conditions (30–33).

Recent studies have hypothesized that as periodontal disease manifests itself as low grade chronic inflammatory condition, it has the potential to contribute to the generation of a systemic inflammatory phenotype (34). This hypothesis is consistent with evidence demonstrating that individuals with periodontal disease present with elevated markers of inflammation (4,35). Notably, these markers are found at lower concentrations among healthy individuals (35). This supports the concept that periodontal disease is an independent factor in the progression of systemic grade inflammation. However, the consideration that periodontal disease may be an independent factor in the risk of diabetes complications is yet to be determined. As seen in figure 1, the difference in survival probabilities among participants self-reporting “good to excellent” and “poor to fair” oral health suggest that there may be specific implications of poor oral health in the oral health systemic pathway. Markedly, the difference in probabilities observed prospectively, suggests that the complications occurring among participants reporting “poor to fair” oral health are chronic in nature. In contrast, participants reporting “good to excellent” oral health were observed having fewer complications with shorter mean follow-up times.

These study results make parallels with clinical studies showing an increase in systemic inflammatory markers in individuals with gingivitis in comparison to those without (36). In particular, studies regarding experimental gingivitis have observed that diabetics develop a greater inflammatory response reacting to experimental plaque accumulation (37). Subsequently, systemic reviews have also found temporary reductions in blood sugar levels among diabetics following non-operative periodontal therapy (4). Although reductions in blood sugar levels were minimally significant just following care, regularly provided periodontal therapy has the potential to improve metabolic control over longer periods of time, thereby potentially altering the inflammatory profile of individuals and reducing the risk of chronic complications. However, the independent nature of periodontal disease in determining the overall risk profile of individuals with chronic disease remains elusive.

The notion that health and health outcomes are socially patterned, may also impact the oral health - diabetes pathway. Although there is considerable evidence regarding the biological mechanism, understanding the quantitative impacts of lifestyles and socially engrained health on the oral systemic link may provide more insight to the role periodontal disease plays in overall health. As such, this concept substantiates the sequential adjustments for social factors and health behaviours in our analysis. Although the social determinants have gained popularity in public health and widespread evidence suggests their impacts are dominant to health behaviours with regards to improving population level health, the impact of health behaviours on diabetes outcomes was notable in our study. Of particular interest was the minimal reduction in the hazard of complications following adjustments for income, education, immigrant status, ethnicity and rural-urban living status. In comparison, adjustments for health behaviours, including physical activity index, alcohol consumption, smoking as well as dental visits, resulted in a significant reduction in the hazard. This may allude to the prominence of health behaviours in a common risk factor approach between oral health and diabetes and may offer guidance to the development of multifaceted interventions that address both social and behavioural factors.

Our study presents with several strengths. Firstly, much of the current literature on oral health and diabetes has focused on community level clinical interventions to assess the association between periodontal disease and diabetes (4). In contrast, our study identifies diabetes complications associated with oral health status at the population level. Secondly, a validated measure of diabetes diagnosis from the Ontario Diabetes Database (ODD) is used, whereas much of the current literature does not specify the validity of a diabetes diagnosis from insurance claims (17,18). Our study also utilizes self-reported oral health status as the exposure variable and proxy for periodontal condition. As self-reported oral health is a multi-faceted measure for overall oral condition, which includes social, psychosocial, economic and cultural components of oral health, it presents as a convenient and effective measure for inferring diabetes complication risk (38). Notably, studies have found that self-reported oral health status is concordant with the clinical need for oral treatment (39,40). The retrospective selection of our cohort and longitudinal follow-up also presents as a significant strength. As oral health status precedes diabetes complications, temporality is achieved. Lastly, as the CCHS has less than 2% missing data and is representative of the Ontario population, our results have generalizability to individuals with diabetes residing in Ontario.

There are a few important limitations to consider as well. Although self-reported oral health is an acceptable proxy for periodontal disease at the population level and for epidemiological surveys, it may not be of adequate agreement to clinical measures of disease (39). Previous studies have shown that self-reported oral health information is highly specific but not as sensitive (41). In the case of periodontal disease, individuals were able to report that they do not have periodontal disease nor did they need dental treatment, with greater accuracy than those who did in fact have the condition or unmet treatment needs (41). Although clinical measures of periodontal disease, such as gingival probing depths, would provide power to our study, such measures have not been linked to electronic medical records in Canada. Having time varying accounts of self-reported oral health status would also strengthen our findings. In regards to our inclusion criteria, although the ODD has a high sensitivity and specificity in capturing diabetic patients from electronic health records, there is still a chance that some individuals might be missing (23). This may include individuals that have not been diagnosed with the condition or those who do not encounter the health care system regularly. However, as the prevalence of undiagnosed diabetes in Canada is approximately 1.13%, missed cases would not significantly impact the results of our study (23). As the periodontal disease-diabetes link is based on the biological mechanism linked to adult onset diabetes, and the ODD does not differentiate between type 1 or type 2 diabetes, our study results may be overestimated. However, as more than 95% of ODD cases are made up of individuals with type 2 diabetes and our study was restricted to adults over the age of 40, the likelihood of overestimation is reduced. Misclassification error could occur for diabetes complications utilized in our study. Although the ICD codes identifying diabetes complications were previously used in other studies, there may be some codes that were missed or others that were misclassified. Lastly, although we were able to control for many potential confounders, there may be residual confounding due to the ambiguities of self-reported data. However, due to the high quality of linked CCHS data and adjustments made for many potential confounders of oral health and diabetes complications, residual confounding was arguably minimized.

In general, our results have both clinical and population level implications for improving overall health. Recent interventions addressing periodontal disease have been driven by collaboration and mostly developed at the community level, including systemic health referrals networks, diabetes screening and oral health education in various settings (42–44). However, a population level intervention or a clear policy addressing systemic health does not exist in Canada. Policies that seek to develop integrated solutions such as collaborative approaches and mixed universal health models should consider the syndemic paradigm of disease in overall health. This unique perspective suggests that as separate entities, periodontal disease and diabetes act synergistically and increase the overall burden of inflammation and disease in the population (45,46). With a distribution of individuals carrying varying burdens of inflammation collaborative approaches for improving overall health are imperative (47). Collaborative strategies should target resources at solutions that address accessibility to those carrying the greatest burden of disease. Inter-professional communication between care givers such as dentists, dental hygienists, primary physicians and specialists should be promoted to enrich a seamless health care experience. While a mixed universal model including dental care is an ideal solution, the private financing and delivery of dental care in Canada remains a challenge. Increasing access to dental care and implementing oral-systemic health assistance programs may offer a solution that mitigates the growing burden of syndemic diseases while managing the current political atmosphere. Identifying providers who are willing to integrate medical and dental care for patients should also be a priority (48,49).

Overall, our findings support the need for interventions that pursue the objective of reducing systemic inflammation. The evidence shows that oral health impacts diabetics and their health outcomes. However, current directions in oral health and diabetes policy provide little indication that this growing concern is being addressed. Policy developments should consider programs that promote access to “whole” health care through integrated health networks that work with physicians, hospital teams and other dental care providers to ensure equitable access and co-ordination for people with treatment needs. Given the growing prevalence of chronic diseases and the impact that oral health status has on the increasing risks of diabetes complications, it is important to develop both targeted and population level strategies in this for reducing overall morbidity and mortality and improving quality of life among Canadians.

